# Chamber-Specific Transcriptomic Insight into Cardiac Development using Guinea Pig and Human Heart Tissue

**DOI:** 10.1101/2025.08.06.666176

**Authors:** Shatha Salameh, Devon Guerrelli, Luther Swift, Anika Haski, Alisa Bruce, Manan Desai, Yves d’Udekem, Nikki Gillum Posnack

## Abstract

The heart undergoes significant molecular and functional adaptations throughout postnatal development. However, to date, our understanding of these dynamic changes in the human heart is limited. Moreover, advances in pediatric cardiac research can be hindered by a lack of preclinical models that accurately reflect human heart maturation. Guinea pigs may serve as a useful model for human cardiac research, as the guinea pig and human myocardium have similar ion channel expression and cardiovascular drug responsiveness. Despite these similarities, gene expression patterns during postnatal heart development have not been comprehensively investigated. In this study, we first characterized transcriptional changes in neonatal, juvenile, and adult guinea pig hearts – identifying gene ontologies and pathways associated with cardiac maturation. Second, we compared the transcriptional profile of right atria and left ventricular tissue to highlight unique and shared chamber-specific patterns in guinea pigs over time. Finally, we conducted a cross-species comparison of the right atrial transcriptome between humans and guinea pigs to identify conserved maturation markers and gene expression patterns. Our findings provide a molecular framework for understanding age- and chamber-specific cardiac development, supporting the guinea pig as a promising preclinical model for studying human heart maturation. By identifying conserved gene programs and developmental markers across species, this study lays the groundwork for age-specific pharmacological strategies and computational models that can help to refine treatment decisions and outcomes for pediatric cardiology patients.

**New and Noteworthy:** Existing knowledge on postnatal heart development and cardiomyocyte maturation is limited. We investigated age-dependent transcriptional changes in neonatal, juvenile, and adult guinea pig hearts - and then conducted a cross-species comparison to identify age-specific patterns that are conserved in the guinea pig and human atria. Expanding our knowledge of chamber- and age-specific gene expression patterns can inform and guide the selection of cardiovascular therapies in the pediatric population, where developmental differences are understudied.

## Introduction

Studies suggest that human cardiomyocytes undergo significant developmental adaptations during the postnatal period, resulting in maturation of myofilaments, the sarcoplasmic reticulum, cardiac ion channel and receptor expression(1–7). However, the depth of our knowledge is limited by the scarcity of human pediatric heart samples, which results in generalized assumptions about developmental temporal patterns due to small sample sizes. Equally important – pediatric human heart tissue samples are collected from patients undergoing palliative or corrected heart surgery, and may not necessarily represent the developmental changes in a normal healthy heart. Experimental animal models can help to address knowledge gaps, but it is important to consider species-specific differences that could hinder the extrapolation of basic research findings to clinically relevant outcomes (8–10). For example, mice and rats differ significantly from humans in developmental heart rate trajectories, myofilament isoform composition, and ion channels/currents that shape the cardiac action potential(11–15). Conversely, guinea pigs (*Cavia porcellus*) share more similarities with humans in terms of action potential shape, ion channel expression, and electrocardiogram metrics(15–17). Additionally, both guinea pigs and humans undergo comparable developmental transitions during gestation, and postnatally both display comparable disease-specific and pharmacological outcomes(16, 18). Further, adult guinea pigs are accepted as a translational cardiac model for drug efficacy and safety testing, due to their high accuracy in predicting human cardiovascular liability to multiple drug candidates(19–21).

Despite select species differences, mammalian models collectively show distinct morphological and physiological phenotypes between the different cardiac chambers (22–25). Further, the electrophysiological and contractile properties of atrial and ventricular cardiomyocytes are intrinsically linked to their distinct developmental origins and unique transcriptional signatures(26–28). Prior work demonstrates regional transcriptional specialization – with ventricular cardiomyocytes prioritizing genes involved in contraction and energy metabolism (22, 29), and atrial cardiomyocytes upregulating genes involved in neurohumoral and endocrine signaling (22). These distinct transcriptional profiles help drive chamber-specific functions that are essential to cardiac performance(22, 27). Notably, these chamber-specific gene expression profiles continue to mature postnatally, providing insight into age-specific adaptations that underlie atrial and ventricular specialization(30).

Although the human and guinea pig share similarities in heart function, the cardiac transcriptome has not been well characterized during postnatal development in either species. To address this gap, we investigated age-dependent transcriptional changes in the neonatal, juvenile, and adult guinea pig heart. Specifically, we identified distinct gene expression patterns and maturation-associated pathways in both the right atrium (RA) and left ventricle (LV) – including chamber-specific and conserved changes throughout development. Finally, we conducted a cross-species comparison of age-specific patterns in the right atrium of humans and guinea pigs. Collectively, our study defines the temporal and spatial transcriptional landscape of the developing guinea pig heart, which can help support its utility as a translational model for better understanding postnatal cardiac maturation.

## Methods

### Human Subjects

Tissue collection and experimental studies were performed in accordance with a Children’s National Hospital Institutional Review Board-approved protocol (IRB# Pro00012146 and STUDY00000198). This minimal risk study involved the collection and preservation of right atrial tissue samples (collected as medical waste), from acyanotic patients who underwent heart surgery at Children’s National Hospital, as previously described (n=160, **Supplemental Table 1**)(31). Tissue samples were designated to one of three age groups: neonate/infant (5-364 days, n=68), children (1-11 years; n=62), and adolescent/adults (12-32 years, n=30). The age groupings were defined prospectively based on designations by the US Department of Health and Human Services, the Food and Drug Administration, and prior evidence supporting the notion that cardiomyocyte maturity is reached after the first decade of life(32, 33).

### Animal model

The Institutional Animal Care and Use Committee of the Children’s National Research Institute approved all animal procedures, which align with the guidelines outlined in the National Institutes of Health’s Guide for the Care and Use of Laboratory Animals and the Animal Welfare Act. Experiments were performed using male and female Dunkin-Harley guinea pigs procured from two suppliers of laboratory animals (Elm Hill Labs: Massachusetts, USA; Charles River Laboratories: Quebec, CA). Animals were housed in conventional acrylic cages within the research animal facility, following standard environmental conditions including a 12 hr light/dark cycle, a temperature range of 18–25°C, and humidity levels maintained between 30 and 70%. To evaluate age and chamber-specific differences, guinea pig tissue (n=30) was categorized into three age groups: neonates (1–2 days old), juveniles (4-10 days old), and adults (>6 months old). Both right atrial (neonates, n=5; juveniles, n=5; adults, n=5) and left ventricular samples (neonates, n=5; juveniles, n=5; adults, n=5) were collected and preserved.

### Gene Expression Analysis

Cardiac tissue samples were submerged in RNAlater stabilization solution (Invitrogen, Waltham, MA, USA) and stored at 4°C for <7 days. RNAlater was subsequently removed and samples were then stored at -80°C until RNA extraction. Total RNA was isolated from right atrial or left ventricular tissue (5-30 mg) and processed using a RNeasy fibrous mini tissue kit with on-column DNase treatment (Qiagen, Germantown MD, USA). RNA concentration was determined using a Qubit assay (Thermo Fisher, Waltham, MA, USA) and RNA quality was assessed using a TapeStation system (Agilent Technologies, Santa Clara CA USA). Only samples of sufficient quality were used in subsequent microarray experiments. 250 ng of total RNA input was primed for the entire length of RNA, including both poly(A) and non-poly(A) mRNA and reverse transcribed to generate sense-stranded targets that were biotin-labeled using a GeneChip WT Plus Reagent kit, and then hybridized to Human Clariom™ S Arrays or GeneChip™ Guinea Pig Gene 1.0 ST Array (Applied Biosystems, Waltham MA USA) for 16 hours at 45°C. After removing the hybridization cocktail, each array was washed and stained on the Fluidics Station F450, and then scanned using an Affymetrix GeneChip Scanner (GCS 3000 7G; Thermo Fisher). Initial quality control data was evaluated using Affymetrix Transcriptome Analysis Console Software. Microarray data were imported and analyzed using the Transcriptome Analysis Console (Applied Biosystems).

### Statistical Analysis

To identify differentially expressed genes (DEGs), data sets were compared between age groups using one-way ANOVA with a p-value threshold <0.05, fold change cutoff of >|1.25|, and a false discovery rate of <0.1. For each DEG, the raw signal intensity data underwent log_10_ transformation and Z-score normalization (Z score = (signal intensity – mean signal intensity)/standard deviation of signal intensity) on a per-gene basis. Z-scores were used to display DEGs as a heatmap using Morpheus(34). Enrichment analysis was performed on a single rank-ordered gene list via analysis tools, including David Bioinformatics, and Enrichr(35–38). Datasets are available via the Gene Expression Omnibus.

## Results

### Postnatal Transcriptional Changes in Guinea Pig Right Atrium and Left Ventricle

To obtain a transcriptomic map of guinea pig hearts, we collected right atrial and left ventricular tissue from three age groups, each comprising five animals. We performed microarray analysis to identify differentially expressed genes (DEGs) and define the biological pathways associated with cardiac maturation within each chamber (**Figure 1**). Notably, principal component analysis (PCA) clearly distinguished between the neonatal, juvenile, and adult age groups in left ventricular tissue. However, in right atrial samples, the neonatal and juvenile age groups presented overlapping boundaries (**Figure 1A, E**). To identify gene ontologies and biological pathways associated with myocardial maturation, we used the Database for Annotation, Visualization, and Integrated Discovery (DAVID) to identify 25 unique annotation clusters in the right atria and 40 clusters in the left ventricle between the neonatal and adult groups (**Figure 1B, F**; **Supplemental Table 2**). Annotation clusters related to maturation in both the RA and LV included those associated with the extracellular matrix (e.g., KW-0272, KW-0084, GO:0062023) and bioenergetics (e.g., KW-0443, cpoc01212, cpoc00071, HSA01100). Using a 1.25-fold expression cut-off and a p-value of 0.05 a total of 2390 DEGs were identified in atrial samples and a total of 3409 DEGs were identified in ventricular samples when comparing neonates versus adults. In right atrial samples, 1106 genes were upregulated and 1284 were downregulated in neonates relative to adults. Similarly, in the left ventricular samples, 1615 genes were upregulated and 1794 were downregulated in neonates relative to adults (**Figure 1C, G**). Using the Enrichr analysis tool, we identified 53/37 gene ontologies that were associated with up/down regulated genes related to biological function in the right atria and 106/6 gene ontologies associated with up/down regulated genes in the left ventricle (**Supplemental Table 2**). A subset of those gene ontologies is shown in **Figure 1D, H**. In both chambers, neonates overexpressed genes associated with the miotic cell cycle process (e.g., *Cdk4, Cdk6, Cdk7*). In the right atrium, neonates underexpressed genes associated with calcium ion transport (e.g., *Casq2, Sln, Gstm2*), cardiac muscle development (e.g., *Bmp10, Mylk3, Sorbs2*, and the ryanodine receptor (e.g., *Camk2d;* **Figure 1D**). In the left ventricle, neonates overexpressed genes associated with glycogen metabolism (e.g., *Igf1, Dyrk2, Irs1*) and muscle organ development (e.g., *Myh6, Itga11, Tcf12;* **Figure 1H**).

**Figure 1.**
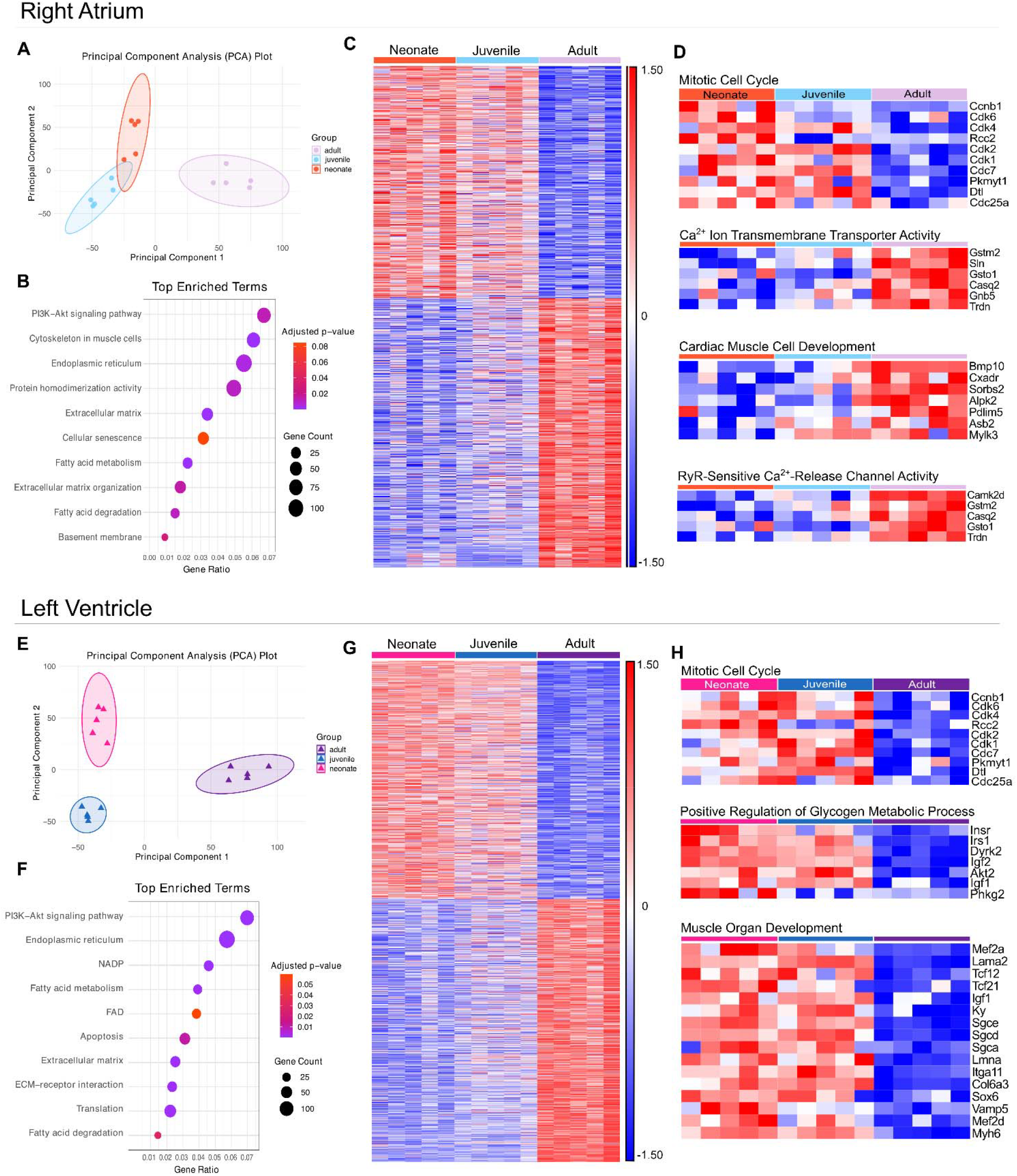
Gene ontology and pathway analysis of differentially expressed genes related to postnatal maturation in guinea pig right atrium (**Top**) and left ventricle (**Bottom**). Principal component analysis between age groups. **(A)** The right atrial variation values were 43.5% (PC1) and 22.6% (PC2), and **(E)** the left ventricular variation values were 43.6% (PC1) and 20.7% (PC2). Subset of annotation clusters (identified using DAVID) that were differentially expressed between neonates and adults in **(B)** right atrial or **(F)** left ventricular samples. Number of genes per cluster is indicated by the dot size, and Bonferroni-adjusted p-value <0.1 by the color. Differentially expressed genes were identified for each chamber between neonatal and adult samples, using one-way ANOVA with a 1.25-fold change cutoff and a 0.1 false discovery rate. A total of **(C)** 2390 DEGs in the right atrium and (G) 3409 DEGs in the left ventricle are shown in the heatmap (each row) for each sample (column); on a per gene basis, the signal intensity was log_10_ transformed and Z-score normalized. **(D,H)** Subset of gene ontologies (identified using Enrichr) that are significantly up- or down-regulated between neonates and adults in right atrial or left ventricular samples. Heatmap shows each gene (row) for each animal (column) within the specified gene ontology; signal intensity was log_10_ transformed and Z-score normalized on a per gene basis. n=5 neonates, n=5 juveniles, n=5 adults.

### Differential and Overlapping Developmental Gene Expression Patterns Between Atria and Ventricles

Next, we compared the transcriptome profiles across both chambers and all three age groups using principal component analysis. Individual age groups (neonatal, juvenile, and adult) clustered distinctly, while atrial and ventricular samples showed overlapping profiles within each age group (**Figure 2A**). Venn diagrams illustrate the differentially expressed genes between right atrial and left ventricular tissue at each developmental stage. A total of 1,309 chamber-specific genes were unique to neonates, 934 were unique to juveniles, and 1,460 were unique to adults. An additional 2,082 genes were shared across all three age groups, representing consistent chamber-specific expression from neonatal through adult stages (**Figure 2B**). Differentially expressed genes between right atrial and left ventricular tissue were identified separately for neonates, juveniles, and adults. Each group of upregulated genes (e.g. upregulated in neonatal atrium, upregulated in juvenile atrium, upregulated in adult atrium) was analyzed using Enrichr to identify enriched biological ontologies (**Supplemental Table 3**). Distinct chamber-specific biological ontologies unique to each age group were identified in either atrial (**Figure 2C**) or ventricular (**Figure 2D**) tissue. In the atrium, biological ontologies that were uniquely overrepresented in juveniles included sodium activity (GO:2000649) while the adult atrium overexpressed calcium handling-related ontologies (GO:0051281, GO:0051591). In the ventricle, biological ontologies that were uniquely overrepresented in neonates included sarcomere organization (GO:0045214) and the fatty acid biosynthetic process (GO:0006633). Unique to the adult ventricle, biological ontologies related to muscle growth and development were overexpressed (GO:0055008, GO:0003229, GO:0003208). Next, we identified chamber-specific biological ontologies shared across all three age groups (**Figure 2E, F**). In the atrium, biological ontologies shared amongst all three age groups included sinoatrial node development (GO:0003163), cardiac conduction system development (GO:0003161), and axon guidance (GO:0007411). Atrial-specific markers conserved across age groups include *Bmp10, Sln*, and *Myl4*. In the ventricle, biological ontologies shared amongst all three age groups included regulation of cardiac muscle cell action potential (GO:0098901), fatty acid beta-oxidation (GO:0006635), and cellular respiration (GO:0045333). Ventricular-specific markers conserved across all three age groups include *Myl2, Myl3*, and *Myh7*.

**Figure 2.**
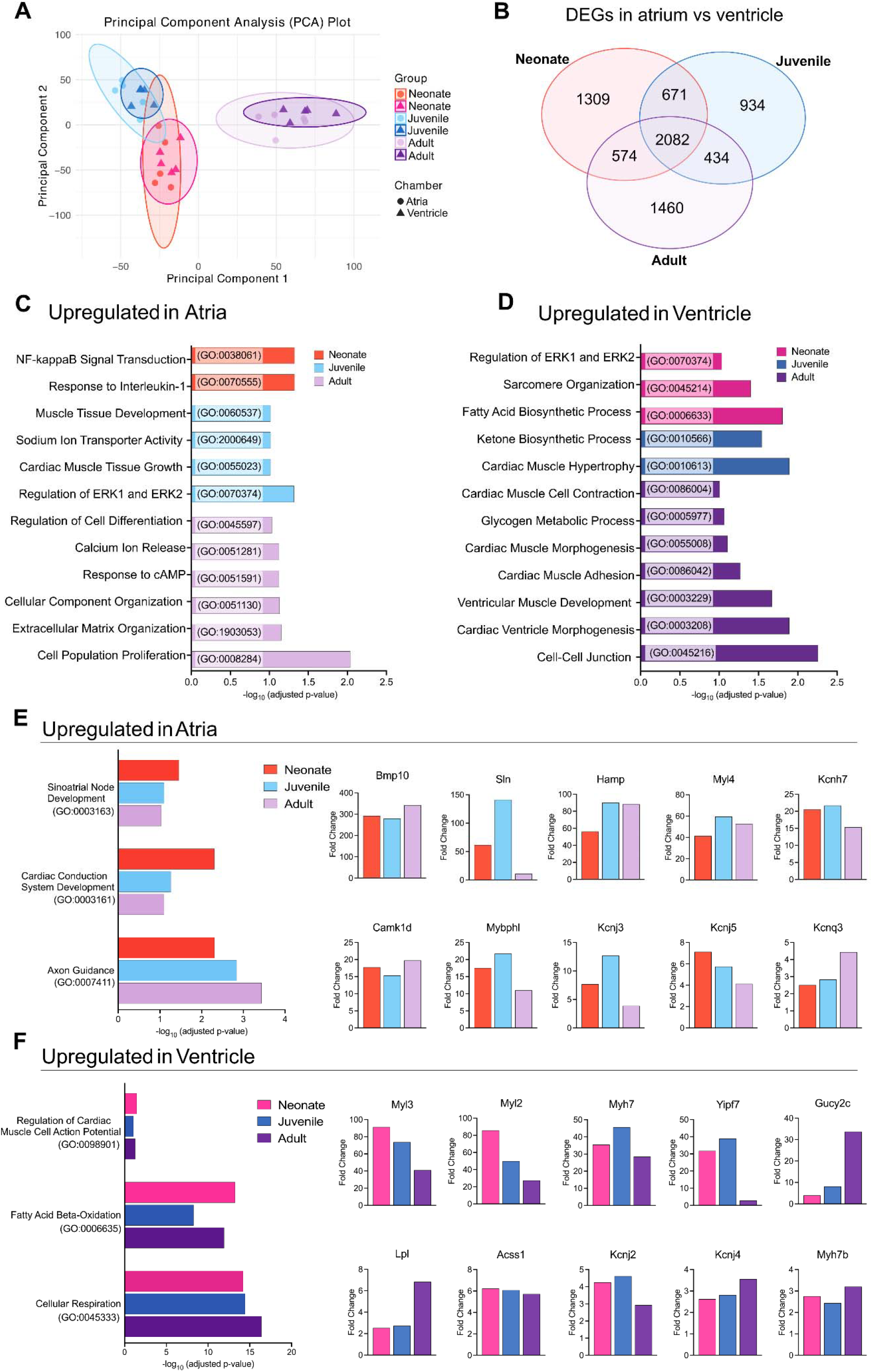
Chamber-specific and conserved gene ontologies across each age group. **(A)** Principal component analysis plot of the transcriptome profiles showing variations between chambers and age groups. The variation values of PC1 and PC2 were 28.3% and 15.6%, respectively. Circles indicate atrial samples and triangles indicate ventricular samples. **(B)** Venn diagrams illustrate differentially expressed genes identified using ANOVA with a 1.25-fold change cutoff and a 0.1 false discovery rate. Upregulated genes in either **(C)** atrial or **(D)** ventricular tissue were used to determine unique annotation clusters for each age groups (identified using Enrichr). Upregulated genes in either **(E)** atrial or **(F)** ventricular tissue were used to identify chamber-specific and conserved annotation clusters (identified using Enrichr). n=5 neonates, n=5 juveniles, n=5 adults.

### Cross-Species Comparison of Human and Guinea Pig Right Atria

Next, we performed a comparative analysis of genes related to postnatal maturation in the human and guinea pig right atrium. Analysis was limited to this chamber due to tissue availability, as a small piece of the right atrium is commonly removed during corrective heart surgery (and later discarded as medical waste). Differential gene expression analysis was performed between the youngest (postnatal days 0–2) and oldest (>6 months) guinea pigs, as well as between the youngest (<1 year) and oldest (>12 years) humans. The resulting gene sets were compared across species using a Venn diagram to identify shared and species-specific transcriptional changes associated with postnatal maturation (**Figure 3A**). Next, we compared gene expression patterns between juvenile (postnatal day 4–10) and adult guinea pigs, as well as children (1–11 years) and adolescent/adult humans. The resulting gene sets were then compared across species to identify shared and distinct transcriptional changes that occur later in the postnatal maturation process (**Figure 3B**). The human and guinea pig hearts shared 667 maturation-related DEGs during early development (**Figure 3A**) and 376 DEGs during later developmental stages (**Figure 3B**). A subset of biological and cellular gene ontologies was identified using Enrichr from the 667 developmental genes shared between humans and guinea pigs (**Supplemental Table 4**). Across species, neonates/infants underexpressed genes associated with fatty acid metabolism (e.g., *PLK3, TPK1*) and the sarcoplasmic reticulum (e.g., *CASQ2, S100A1, SLN*). The differential expression of genes related to the positive regulation of signal transduction (e.g., *LAMC1, LAMA2, DACT1*) occurred with age in both species (**Figure 3C**). Scatter plots display the fold changes of overlapping upregulated genes in the youngest age group in both guinea pigs and humans. Bar graphs display the fold changes of selected genes of interest, including cell-cycle-related genes (e.g., *CDC7, TOP2A, MKI67*), which represent conserved markers of immature cardiomyocytes across species during early development (**Figure 3D**). In contrast, scatter plots show the fold changes of overlapping genes upregulated in later development and humans (**Figure 3E**). Bar graphs display the fold changes of selected genes of interest, including calcium-handling genes (*S100A1, SLN, CASQ2*) and genes encoding ion channels (*KCNJ3, KCNA3*), which represent conserved markers of cardiomyocyte maturation across species during later postnatal development.

**Figure 3.**
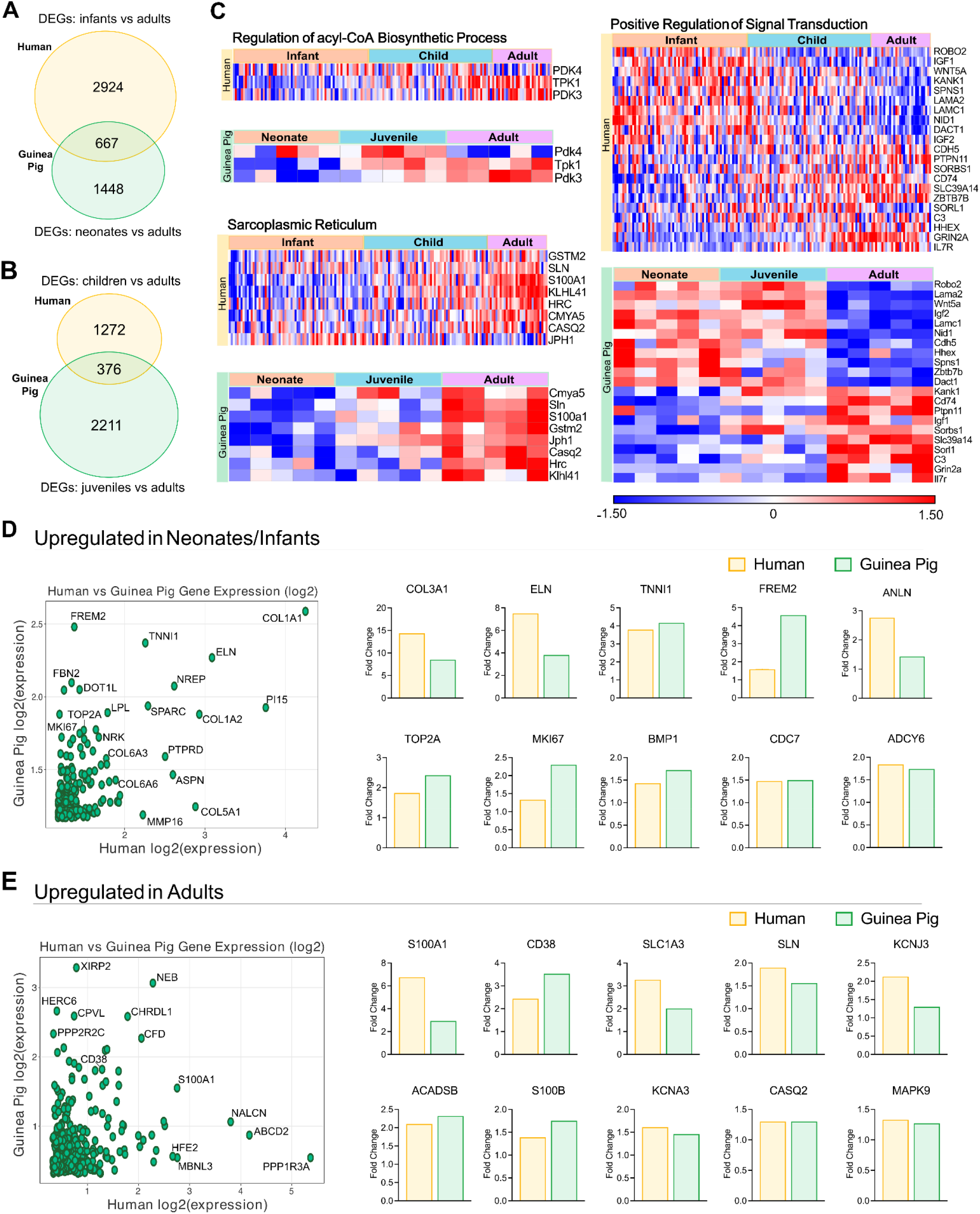
Developmental species comparison using human and guinea pig right atrial samples. **(A, B)** Venn diagrams show differentially expressed genes between species, comparing either the youngest or middle age group to adults. **(C)** Subset of shared gene ontologies (identified using Enrichr) from genes differentially expressed between the youngest age group and adults (Note: humans identified in yellow, guinea pigs in green). Scatter plots show the fold change of genes **(D)** upregulated in the youngest age group relative to adults or **(E)** upregulated in adults relative to the youngest age group. For scatter plots, only conserved genes in both humans and guinea pigs are shown. The accompanying bar graphs highlight selected genes of interest. Humans: n=68 neonates/infants, n=62 children, n=30 adolescent/adults. Guinea pigs: n= 5 neonates, n=5 juveniles, n=5 adults.

### Conserved Temporal Expression Patterns in Right Atrial Maturation Across Species

Finally, we analyzed the expression trajectories of shared genes across the three postnatal age groups in humans and guinea pigs. This enabled direct comparison of the temporal dynamics across two time periods (early development: infant/neonate to child/juvenile; late development: child/juvenile to adolescent/adults) of conserved atrial maturation markers between species (**Figure 4**). Immature isoforms of contractile proteins (TNNI1) and calcium signaling modulators (ADCY6) were upregulated in neonates/infants relative to adults in both species. Genes related to calcium-handling (*SLN, CASQ2*), energy metabolism (CKM) and smooth muscle function (ACTA2) consistently increased in expression from early to later stages of development. Genes encoding the calcium binding protein (S100A1) and myosin light chain 4 (*MYL4*) showed the most pronounced increase in expression during the later developmental period. In contrast, genes encoding caveolae formation (*SDPR*) exhibited the most drastic increase in expression during the early developmental period. Interestingly, although ultimately upregulated in adults across species, a subset of genes (TAGLN, CAV1, SLN) had more gradual increases in expression during early postnatal stages in either humans and guinea pigs (**Figure 4A**). These temporal patterns are important to consider when selecting atrial maturation markers, as their expression dynamics may vary depending on the specific developmental timepoints examined.

**Figure 4.**
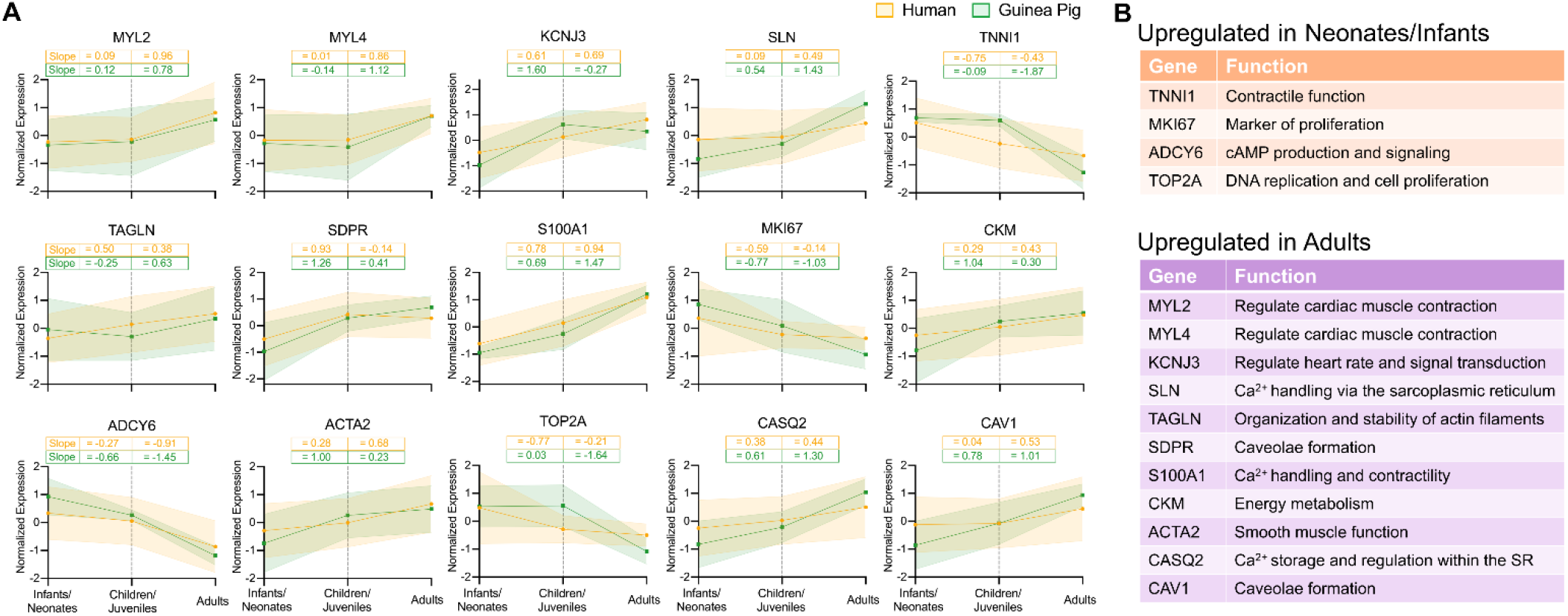
Atrial markers that are conserved across species. **(A)** Line plots show the expression pattern of right atrial-specific markers temporally across three age groups in both humans and guinea pigs. Signal intensity was log_10_ transformed and Z-score normalized on a per gene basis, and the slope was calculated between each consecutive age group. **(B)** Functional descriptions of atrial markers highlight their biological roles and relevance to cardiac development. Humans: n=68 neonates/infants, n=62 children, n=30 adolescent/adults. Guinea pigs: n= 5 neonates, n=5 juveniles, n=5 adults.

## Discussion

To our knowledge, this is the first transcriptomic comparison between human and guinea pig heart tissue. First, our data showed miotic cell cycle markers (e.g. *TOP2A, CDC7, MKI67*) are overexpressed early in development, regardless of chamber or species. This is expected, as cardiomyocyte maturation involves an age-dependent transition from hyperplasia to hypertrophic growth. Prior work in human tissue suggests that cell cycle gene expression declines after the first few months of life, with cardiomyocyte proliferation largely restricted to the neonatal period(39–41). Second, our study revealed that humans and guinea pigs share age-dependent gene expression patterns, including an enrichment of key ontologies related to fatty acid metabolism and sarcoplasmic reticulum function – highlighting conserved postnatal shifts in metabolism and calcium-handling processes. Third, we reported unique chamber-specific gene ontologies that were enriched at each developmental stage – including atrial upregulation of ontologies related to conduction and signal propagation, and ventricular upregulation of ontologies related to action potential regulation and bioenergetics. Principal component analysis revealed unique temporal patterns of maturation between the chambers, as ventricular samples yielded greater separation between age groups compared to atrial samples. This aligns with prior work noting distinct differentiation pathways between atrial and ventricular cardiomyocytes(42, 43).

While much of the existing literature has focused on ventricular cardiomyocyte maturation, our study highlights distinct transcriptional markers that define atrial maturation and reveal regulatory programs essential to atrial identity(44). For example, *SLN*, which encodes the endogenous calcium cycling regulator sarcolipin, was significantly upregulated in atrial tissue of adolescent/adult humans and adult guinea pigs in this study(45–47). These findings are in agreement with previous reports documenting that sarcolipin expression increases with development in the atria, but remains undetectable in the ventricle(48, 49). We also found that *KCNJ3*, which interacts with *KCNJ5* to form the acetylcholine-activated potassium channel (responsible for I_KAch_) was upregulated in the right atria relative to the left ventricle, and its expression was increased in adults in both species(50, 51). This is supported by previous reports showing that *KCNJ3* is atrial-specific and constitutively active in chronic atrial fibrillation, suggesting a role in both electrophysiological maturation and susceptibility to arrhythmogenic remodeling(52–55). Additional experiments in guinea pigs and human tissue support our findings, wherein *KCNJ3* expression was most abundant in atrial tissue(56–58).

Despite their prevalence in cardiac research, small rodents such as mice and rats are physiologically distinct from humans and translation of their genetic findings requires much caution(14). For example, a well-established species difference is the opposite developmental regulation of *Myh6* and *Myh7* (genes encoding alpha and beta isoforms of myosin heavy chain), whereby the faster heart rate in adult rodents is supported by expression of *Myh6* isoform with higher ATPase activity and contraction/relaxation kinetics(12, 59, 60). Conversely, in our study, we reported that guinea pigs are developmentally similar to humans – with an age dependent decrease in *Myh6* and increase in *Myh7* in the ventricle. We also detected increased *Myl2* and *Myl3* (genes encoding myosin light chain isoforms) in ventricular tissue, which reportedly corresponds with chamber-specific differences genes(61). Additionally, we reported similar developmental trends in calcium-handling genes (e.g. *CASQ2, S100A1, SLN*) between human and guinea pig atrial tissue samples.

In summary, our study provides new insight into conserved molecular programs of cardiac maturation and also supports the notion that guinea pigs can serve as a preclinical model to better understand postnatal heart development. Expanding our knowledge of chamber- and age-specific gene expression patterns can inform and guide the selection of cardiovascular therapies in the pediatric population, where developmental differences in signaling pathways, ion channel and receptor expression are understudied. By capturing transcriptional changes across species and developmental stages, our findings can serve as a molecular framework for future work on cardiac physiology in the immature heart.

## Limitations

First, our transcriptomic analysis utilized microarray analysis, which does not capture cell-type–specific gene expression changes. Future studies using single-cell or single-nuclei approaches could refine our understanding of how individual cell populations mature over time, taking into account the cellular heterogeneity of the heart(62). Second, while the guinea pig shares many physiological and pharmacological similarities with humans, interspecies differences in developmental timing, lifespan, and gene regulation may affect the interpretation of conserved gene expression patterns. Third, we derived our human data from pediatric patients with acyanotic congenital heart disease, which may/may not represent the developmental timeline of the “healthy” human heart. Additionally, the cross-species comparison was limited to the right atrium due to the limited availability of tissue specimens from pediatric patients. As such, cross-species maturation patterns across the other cardiac chambers were not included. Finally, our analysis infers biological processes from gene expression alone, without parallel protein or electrophysiological data. Future studies integrating proteomics and functional assays will be essential to link these molecular and physiological outcomes during cardiac maturation across both humans and guinea pigs.

## Acknowledgements

We gratefully acknowledge Susan Knoblach, Karuna Panchapakesan, and the Children’s National Research Institute Genomics and Bioinformatics Core for assistance with microarray experiments. We also acknowledge additional members of the cardiac surgery operating room team, including Aybala Tongut, Blezzy Bote, Alecia Byrd, Jannette Callos, Lula Curry, Kenisha Cyrus, Moozhda Hanif, Evelyn Ravizee, Sandra Sunderland, and Hyung Mi (Grace) Yang for their assistance with tissue sample procurement and coordinators who assisted with IRB protocols and the consenting process, including Alix Fetch, Alyssia Venna, Desiree Nwanze, and Carlos Carhaus.

## Author Contributions

DG, AB, MD, YD screened acyanotic CHD patients for enrollment and preserved atrial tissue samples during cardiac surgery for this biomedical research study. SS, DG, and NGP conceived and designed the study. SS, DG, AH, LS, NGP performed experiments, analyzed and interpreted data. SS prepared figures, SS and DG prepared tables. SS and NGP drafted the manuscript. All authors revised the manuscript or provided critical intellectual content, and all authors approved the final manuscript.

## Data Availability

Derived data supporting the findings of this study are available from the corresponding author (NGP) upon request. Datasets are available via the Gene Expression Omnibus. Supplemental Files are available at https://figshare.com/s/790d24d9b0299332b22f

